# Cell state-dependent chromatin targeting in NUT carcinoma

**DOI:** 10.1101/2023.04.18.537367

**Authors:** Artyom A. Alekseyenko, Barry M. Zee, Zuzer Dhoondia, Hyuckjoon Kang, Jessica L. Makofske, Mitzi I. Kuroda

## Abstract

Aberrant transcriptional programming and chromatin dysregulation are common to most cancers. Whether by deranged cell signaling or environmental insult, the resulting oncogenic phenotype is typically manifested in transcriptional changes characteristic of undifferentiated cell growth. Here we analyze targeting of an oncogenic fusion protein, BRD4-NUT, composed of two normally independent chromatin regulators. The fusion causes the formation of large hyperacetylated genomic regions or megadomains, mis-regulation of *c-MYC*, and an aggressive carcinoma of squamous cell origin. Our previous work revealed largely distinct megadomain locations in different NUT carcinoma patient cell lines. To assess whether this was due to variations in individual genome sequences or epigenetic cell state, we expressed BRD4-NUT in a human stem cell model and found that megadomains formed in dissimilar patterns when comparing cells in the pluripotent state with the same cell line following induction along a mesodermal lineage. Thus, our work implicates initial cell state as the critical factor in the locations of BRD4-NUT megadomains. These results, together with our analysis of c-MYC protein-protein interactions in a patient cell line, are consistent with a cascade of chromatin misregulation underlying NUT carcinoma.

## INTRODUCTION

NUT carcinoma (NC) arises from squamous cells typically lining internal organs. The defining feature of NC is rearrangement of the *NUTM1* gene (French et al. 2003, French 2018). *NUTM1* expression is normally restricted to germ cells, but as a result of chromosomal translocation, NUTM1 becomes the ’ partner of a widely expressed fusion protein. Somatic expression of NUTM1 protein is problematic because its normal function is to hyperacetylate histones for their removal during spermatogenesis (Shiota et al. 2018, Rousseaux et al. 2022). Thus, it is a powerful attractor and facilitator of the p300/CBP histone acetyltransferases (Wang and You 2015, Ibrahim et al. 2022, Yu et al. 2023).

The 5’ translocation partner of NUTM1 fusions in NC provides broad expression and chromatin-association (French 2018). The most common 5’ partners are bromodomain containing proteins such as BRD4, that bind acetylated histones (Zeng and Zhou 2002). BRD4 is a co-activator important for expression of many key genes including *c-MYC* (Filippakopoulos et al. 2010, Kotekar et al. 2022). The BRD4-NUT fusion is a powerful oncoprotein which forms distinctive nuclear foci (French 2018). We previously found that at the genomic level, these foci correspond to ∼200, hyperacetylated expanses of chromatin that reach up to 2 Mb in size (Alekseyenko et al. 2015a, Eagen and French 2021). Based on their unprecedented size, we named these BRD4-NUT-associated regions “megadomains.”

Megadomains can be induced in non-NC cells via transgenic expression of the BRD4-NUT fusion protein. Interestingly, within hours of induction in HEK293T cells, megadomain formation typically starts at a subset of enhancers, and subsequently spreads acetylation long distances in cis, sometimes filling whole topologically associated domains (TADs) (Alekseyenko et al. 2015a). Spreading is thought to occur when the BRD4 bromodomain binds acetylated nucleosomes and the NUTM1 portion of the fusion protein attracts the EP300 lysine acetyltransferase, resulting in a self-perpetuating, feed-forward loop (**Fig. 1A**).

**Figure 1.**
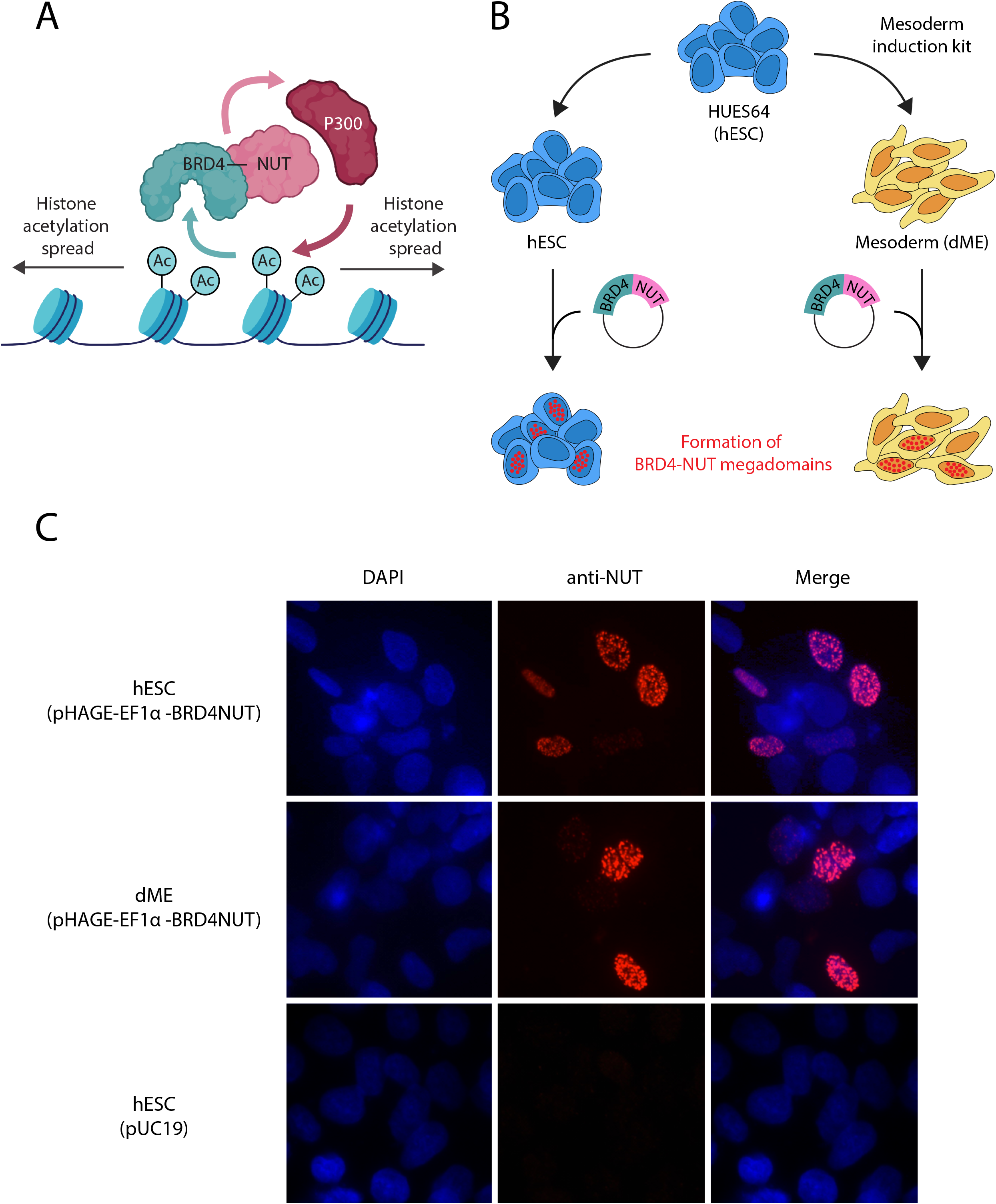
Induction of megadomains in HUES64 human embryonic stem cells. A. Model for formation of BRD4-NUT megadomains. B. Strategy for comparison of megadomain induction following transfection of HUES64 cells maintained in a pluripotent state or after differentiation along a mesodermal lineage. C. Anti-NUT immunostaining after transient transfection of HUES64 cells with a BRD4-NUT expression plasmid or a pUC19 negative control (400x magnification). Formation of NUT nuclear foci is dependent on expression of the BRD4-NUT transgene.

Megadomain locations, mapped by ChIP-seq of NUT or H3K27 acetylation, are strikingly distinct in different NC patient cell lines, with the notable exception of the extensive regulatory region of the *c-MYC* gene which harbors a megadomain in all patient cell lines examined to date (Alekseyenko et al. 2015a, Alekseyenko et al. 2017). In support of a model in which megadomains drive NC, treatment of NC patient cells in culture with a small molecule, JQ1, that inhibits interaction of the BRD4 bromodomains with acetylated lysines, leads to loss of megadomains, down-regulation of *c-MYC*, and differentiation of NC cells (Filippakopoulos et al. 2010, Alekseyenko et al. 2015a). The critical importance of *MYC* was revealed in studies in which ectopic *c-MYC* transgene expression reversed the differentiation induction caused by BRD4-NUT knockdown (Grayson et al. 2014). Furthermore, non-BRD4-NUT-dependent pathways such as *KLF4* expression help maintain *c-MYC* levels in cells resistant to JQ1 (Liao et al. 2018). Thus, understanding the targeting mechanism of megadomains, as well as the molecular interactions of c-MYC protein in NC cells should provide valuable insights into this therapeutically intractable disease.

Here we report our investigation into three questions relevant to this unusual, chromatin-driven disease. First, are the differences in megadomain locations in different patients due to differences in genotypes or instead related to their tissues of origin or epigenetic state? Second, is *c-MYC* the common target in NC patients because there is a powerful growth selection for cells mis-expressing this oncogene, or is *c-MYC* targeting somehow intrinsic to BRD4-NUT across cell states? Third, since MYC proteins are generally considered undruggable (Dang et al. 2017), can identification of c-MYC protein-protein interactions in an NC patient cell line reveal critical co-factors for therapeutic intervention?

RESULTS AND DISCUSSION

### BRD4-NUT megadomain locations are determined by cell state

Our primary goal was to ask why megadomain patterns were distinct in different cancer patient cell lines. From our previous work, we already knew that this was unlikely to be due to a random, stochastic process, as megadomain patterns were remarkably reproducible in replicates induced in cell lines with no prior exposure to BRD4-NUT after transfection of a BRD4-NUT transgene (Alekseyenko et al. 2015a). Therefore, we proposed two hypotheses to account for this distinctive targeting: i) Different patients’ cells have critical DNA sequence variations, which could explain the divergence of megadomain patterns or ii) Different transcriptional or epigenetic states are the key to megadomain pattern formation. For example, NC can arise in squamous cells lining the lung or salivary gland, so the patterns might reflect these initial tissue-of-origin differences.

To distinguish between these hypotheses, we selected a human stem cell model to study how BRD4-NUT megadomain formation responds to changes in cell state (**Fig. 1B**). We purchased HUES64 cells (Chen et al. 2009) from the Harvard Stem Cell Institute iPS core (https://ipscore.hsci.harvard.edu/home) and maintained them in the pluripotent state on matrigel in mTeSR1 media (see Methods). We chose the mesodermal lineage as our comparison cell state because a parallel study confirmed that treatment of HUES64 cells with a commercially available kit (Stemcell Technologies) produced conversion to ∼98% EpCAM-negative/NCAM-positive mesodermal cells by FACS analysis (Naxerova et al. 2021). To express the fusion oncoprotein in hESC and in cells differentiated along a mesodermal lineage (dME), we constructed pHAGE-EF1α -BRD4NUT, in which the fusion cDNA was driven by a strong constitutive promoter derived from the human elongation factor EF-1α gene (**Datafile S1**).

Following transient transfection, we performed immunostaining to test whether expression of BRD4-NUT protein caused the formation of NUT nuclear foci in the two distinct conditions. In each case, we found that foci qualitatively formed in similar number and appearance to those of endogenous BRD4-NUT in cultured patient-derived cells (**Fig 1C**). Nucleofection efficiency of the DNA into hESC was typically around 30%. Since the NUT epitope was uniquely expressed in successfully transfected cells, this allowed our experiments to proceed without drug selection or cell-sorting.

Encouraged by the appearance of nuclear foci after transfection, we performed ChIP-seq using anti-NUT antibodies (Cell Signaling). We scaled up the Lipofectamine 3000 transfection protocol to obtain 5×10^7^ hESC or dME for each ChIP experiment, and performed two independent biological replicates for each cell state. In each experiment BRD4-NUT was expressed for 24 hours. Our results confirmed that BRD4-NUT produced megadomains in highly reproducible patterns in replicates of the same cell state (**Fig 2AB**). In contrast, we found that megadomains were formed in highly divergent patterns when cells in the pluripotent state were compared to the same cell line following induction along a mesodermal lineage **(Fig 2A-D, Datafile S2)**. We conclude that BRD4-NUT initiates megadomain formation in response to the pre-existing transcriptional or epigenetic state of recipient cells.

**Figure 2.**
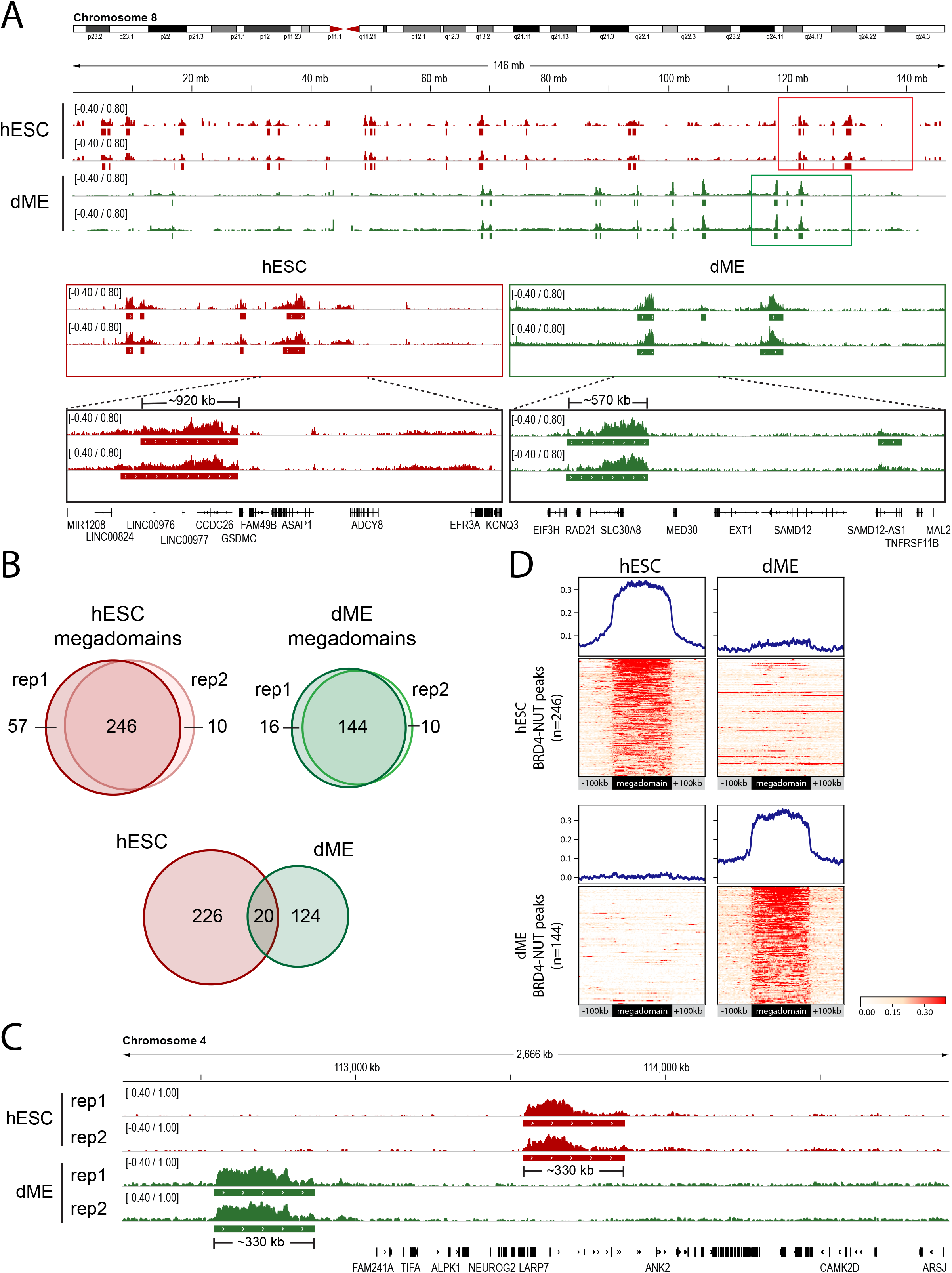
Comparison of BRD4-NUT megadomains by ChIP-seq. A. Top: Human chromosome 8 IGV browser view of log_2_ IP/Input BRD4-NUT read counts from ChIP-seq experiments performed in HUES64 (red) and derived mesoderm dME (green). Middle: Boxed regions are enlarged, showing the strong concordance between independent biological replicates. Bottom: Examples of cell type-specific megadomains in hESC and dME are indicated along with their respective genomic sizes. Transcription units are displayed below the ChIP-seq profiles. B. Venn diagrams showing the strong reproducibility of independent biological replicates of the same cell state, and the divergence when comparing the same cell line in different cell states. C. IGV browser view comparing differences between cell states in a 3 Mb region of chromosome 4. D. Metagene profiles and heatmaps of BRD4-NUT megadomains calculated as log2 ChIP/Input. The x-axis shows megadomains scaled to the average size of 180 Kb, flanked by linear 100 Kb segments of upstream and downstream sequences. Top: Each row of the heatmap represents a megadomain sorted in descending order for BRD4-NUT occupancy in HUES64, revealing many HUES64-specific megadomains (left) along with a few that overlap with dME megadomains (right). Bottom: Heat map of dME-specific megadomains sorted in descending order (right) with the corresponding regions of HUES64 (right).

### Megadomains form at N-MYC rather than c-MYC in pluripotent cells

We next asked whether the *c-MYC* regulatory region, harboring a common megadomain in all NC patients studied to date (Alekseyenko et al. 2015a), was a target of BRD4-NUT in HUES64 stem cells and found that it was not (**Fig 3A**). However, the *MYC* family of cellular oncogenes is comprised of three related genes, *c-MYC*, commonly known as *MYC*, as well as *N-MYC* and *L-MYC* (Henriksson and Luscher 1996, Facchini and Penn 1998). While the *c-MYC* regulatory region was not a target, we found that a prominent megadomain formed on the *N-MYC* regulatory region in HUES64 pluripotent cells (**Fig 3A**). To test whether this might be common to other pluripotent cell lines we transfected Ntera-2 cells, a human embryonal carcinoma cell line (Lee and Andrews 1986), and found similar results, ie. the *N-MYC* regulatory region was a prominent target of BRD4-NUT megadomain formation (**Fig 3B**). As these experiments were performed after 24 hours of transgene expression, selection for BRD4-NUT-driven proliferation was unlikely to be a factor in this association. *c-MYC* and *N-MYC* have similar functions but distinct expression patterns during development (Malynn et al. 2000). Therefore, we speculate that BRD4-NUT may have intrinsic attraction to the relevant *MYC* gene in undifferentiated cell types, potentially because endogenous BRD4 is normally a key co-activator of *MYC* during proliferation.

**Figure 3.**
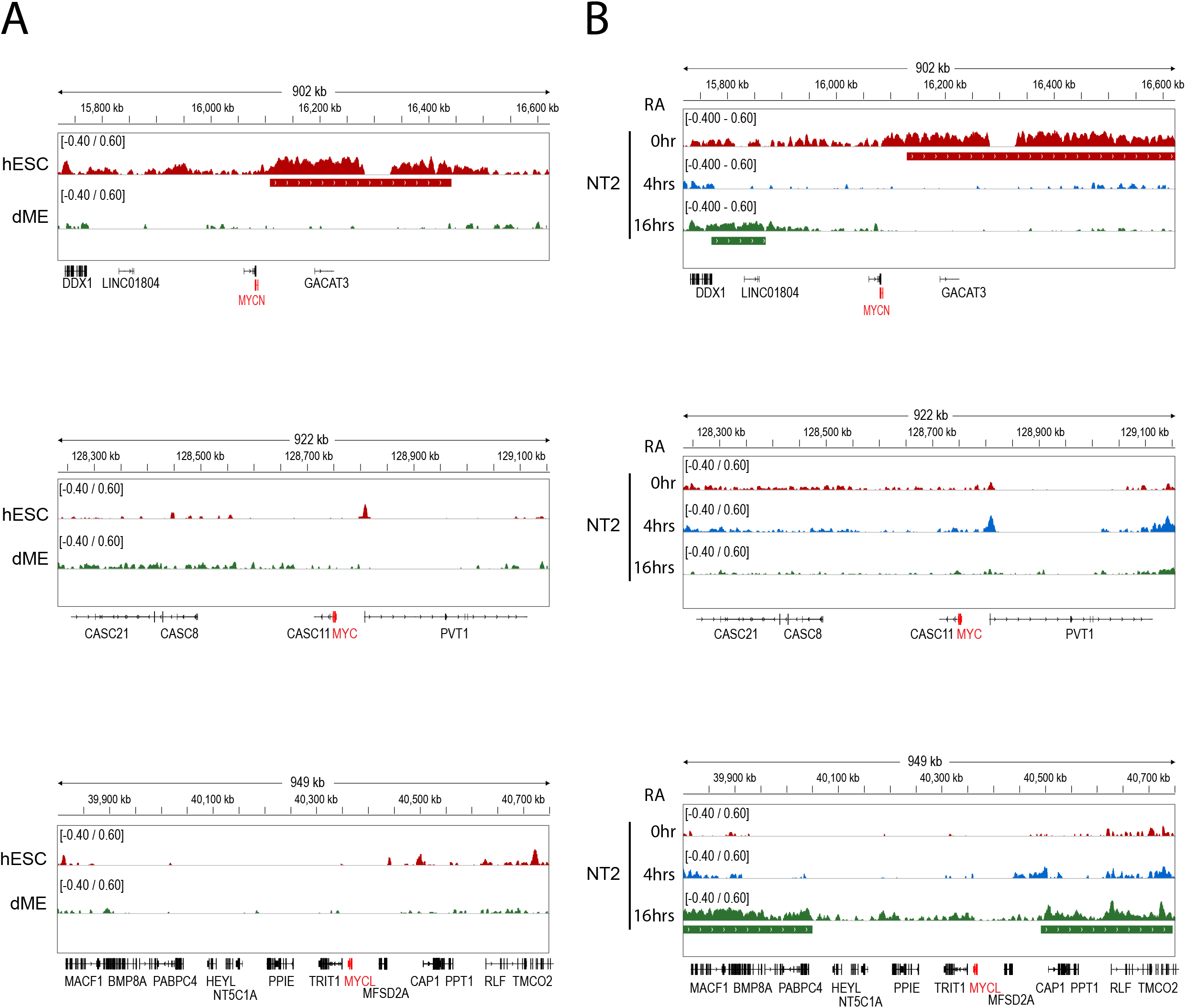
Comparison of BRD4-NUT ChIP-seq of the *N-MYC, c-MYC* and *L-MYC* loci. *N-MYC, c-MYC*, and *L-MYC* IGV browser views of log_2_ IP/Input BRD4-NUT read counts from ChIP-seq experiments performed in A. HUES64 and derived mesoderm dME, and B. Ntera2 cells at 0, 4, and 16 hours of retinoic acid induction, which causes differentiation toward a neural lineage. Both *N-* and *c-MYC* have large adjacent regulatory regions encompassing non-coding RNA transcripts (such as *GACAT3* and *PVT1*) but lacking additional protein-coding genes, while *L-MYC* resides in a region with many protein-coding genes. Colored bars below the ChIP profiles indicate megadomains (>96 Kb contiguous occupancy of BRD4-NUT).

The devastating effect of MYC over-expression is well established in human cancer, but less is known regarding whether dysregulated MYC might acquire aberrant or unexpected protein interactions when driving malignancies. Therefore, we next turned to analysis of the downstream consequences of BRD4-NUT-driven *c-MYC* dysregulation in an NC patient cell line at the level of protein-protein interactions.

### c-MYC protein interacts with the NuA4 acetyltransferase complex in NC797 cells

Given the strong evidence for dysregulation of *c-MYC* as the common feature of NC, we turned our focus to its protein interactions in a well-characterized NC cell line, NC797 (Toretsky et al. 2003). For many years comprehensive proteomic analyses of MYC interactions were impeded by technical challenges. In addition to a relatively short half-life, it has been difficult to extract intact MYC complexes from chromatin using traditional biochemical methods. This has been overcome by the use of biotinylation as a high affinity tag (Kim et al. 2010) or by the implementation of BioID (Roux et al. 2018), in which proximity labeling alleviates the need for release of MYC protein from chromatin to identify interactors (Dingar et al. 2015, Kalkat et al. 2018). Using BioID, the Penn laboratory has mapped the interactions of both the NuA4 (KAT5/TIP60) and STAGA (KAT2A/GCN5) complexes to c-MYC homology box II (MBII). Furthermore, they discovered that MBII is linked to both complexes through interaction with TRRAP and confirmed the importance of MBII in stimulation of acetylation by c-MYC (Kalkat et al. 2018). Thus, comprehensive proteomics has strongly confirmed the initial discoveries of c-MYC association with acetyltransferases (McMahon et al. 2000, Bouchard et al. 2001, Frank et al. 2001, Frank et al. 2003, Liu et al. 2003).

Here, we applied an orthogonal approach, BioTAP-XL (Alekseyenko et al. 2015b), to characterize c-MYC protein interactions in NC and non-NC cells without requiring initial release from chromatin. BioTAP-XL employs crosslinking, sonication, stringent two-step affinity purification, and analysis of enrichment over input by mass spectrometry. Using BioTAP-XL, we found that c-MYC interacts primarily with subunits of the NuA4 acetyltransferase complex, but not STAGA in NC797 cells (**Fig. 4A**). To test whether the lack of STAGA enrichment might be related to our crosslinking method rather than the NC cell type, we performed the BioTAP-XL experiment in the non-NC cell line HEK293T, which was used in the previous BioID studies (Kalkat et al. 2018). Similar to NC797 cells, we found enrichment of NuA4 components with BioTAP-MYC in HEK293-TREx cells, but we also observed enrichment of STAGA components such as ATXN7 and KAT2A (**Fig 4B**). Therefore, c-MYC interaction with NuA4 but not STAGA appears to be a characteristic of the NC797 patient cell line rather than somehow biased by our crosslinking approach, although it should be noted that BioTAP-tagged c-MYC was expressed from its endogenous locus in NC797 cells, and ectopically expressed in HEK293T cells.

**Figure 4.**
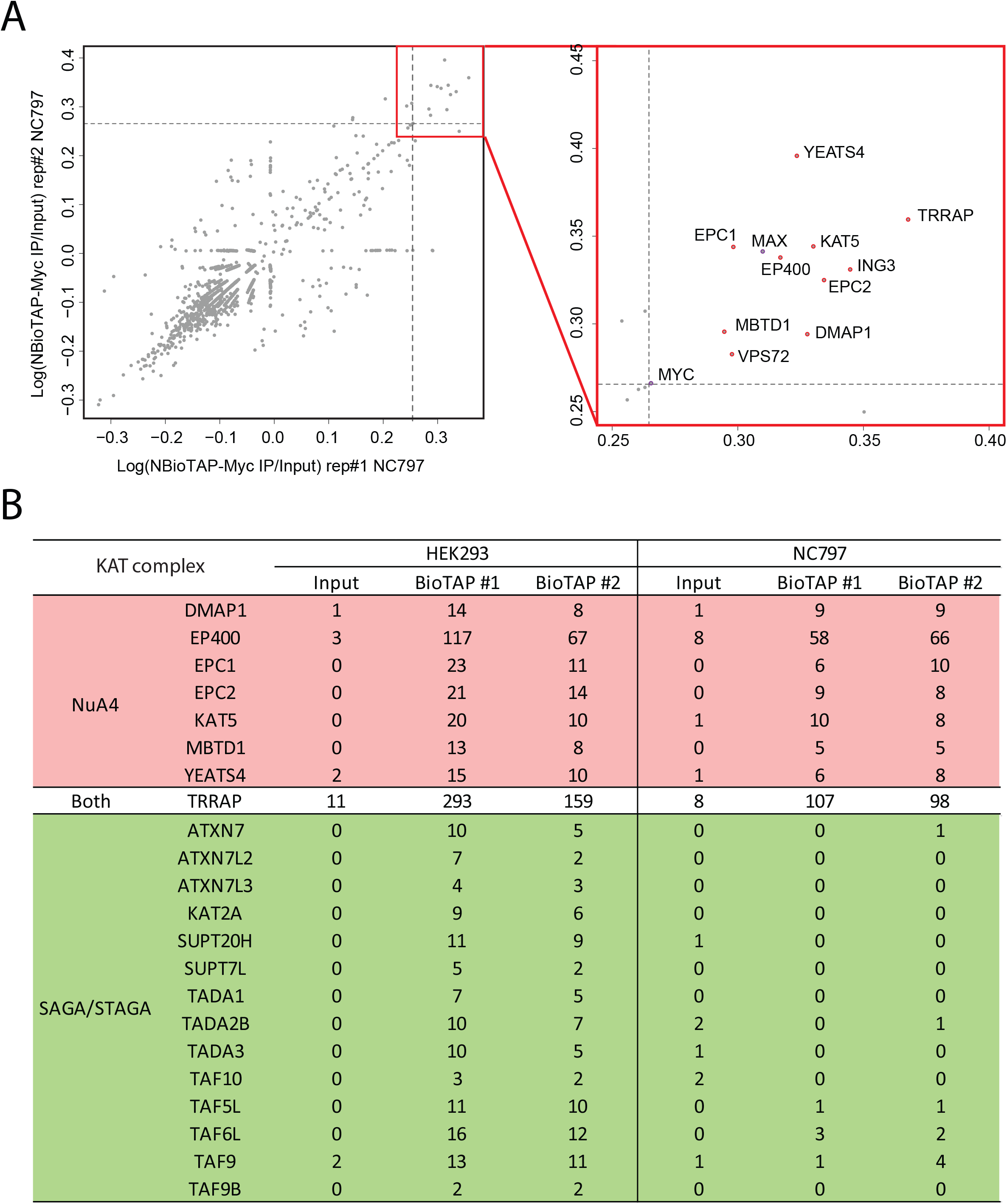
MYC protein interactions in NC797 cells. A. Scatterplot depicting enrichment over input of proteins from N-terminally tagged c-MYC using BioTAP-XL affinity purification in NC797 cells. Each gray dot represents an individually identified protein, where its position is a measure of enrichment efficiency based on the number of total peptides identified in pulldown compared to input and normalized to the molecular weight of the protein. The dashed gray lines denote the 99^th^ percentile of enrichment. The right plot is a zoomed in region of the most enriched proteins in the left plot from replicate experiments. B. Comparison of the total peptides mapping to NuA4 and STAGA components in HEK293T and NC797 cells following BioTAP-XL of c-MYC.

As chromatin acetylation complexes were the most enriched MYC interactors (top 1%) using our stringent crosslinking, tandem affinity purification approach, our results support the extensive previous work cited above, in which acetyltransferase activity is strongly linked to the biological function of c-MYC. Previous nascent RNA-seq analyses revealed transcription of the STAGA acetyltransferase *KAT2A* gene at similar levels in HEK293T and NC797 cells (**Supp Fig S1**). Thus, it remains to be determined why the NC cells we studied here show strong enrichment for MYC interaction with NuA4 but not STAGA. Demonstrations that c-MYC is dependent on acetyltransferases for its function have suggested potential therapeutic options to what is generally considered an undruggable oncoprotein (Mustachio et al. 2020). Our results suggest that development of specific inhibitors of KAT5/TIP60 for synergy with JQ1 could be particularly relevant to the treatment of NC.

In conclusion, while much remains to be discovered regarding BRD4-NUT chromatin-driven oncogenesis, it is striking that its targeting is clearly related to cell state rather than to genotype or DNA sequence. Although we did not directly assess acetylation in the current study, we speculate that differential targeting of BRD4-NUT is likely due to the underlying differences in pre-existing genomic acetylation patterns in pluripotency vs differentiation, based on our prior work mapping the initiation of megadomain formation to a subset of enhancers in HEK293T cells (Alekseyenko et al. 2015a). Thus, acetylation is implicated in both the initiation of megadomains and their aberrant transcriptional consequences. c-MYC up-regulation and strong physical association with acetyltransferases confirms a third connection to a cascade of aberrant acetylation underlying NUT carcinoma.

## MATERIALS AND METHODS

### Cell culture

We purchased HUES64 from the Harvard Stem Cell Institute iPS core facility and maintained them on matrigel (Corning) in mTeSR1 media (Stemcell Technologies). Cells were routinely passaged as clumps after dissociation with 10uM EDTA for 3 minutes as needed. For mesoderm formation, single hESC cells were plated at a density of approximately 5×10^5^ per each well of a 6 well plate in the presence of ROCK inhibitor and exposed to commercially available mesoderm induction media (Stemcell Technologies 05221).

### BRD4-NUT expression in hESC and mesodermal lineage cells

#### Transfection of hESC

7×10^6^ cells were dissociated into single cells with Accutase (Thermo Fisher) at 37°C for 8 -10 minutes. 10ml of mTeSR1 media with 10 uM ROCK inhibitor were added and cells were centrifuged 200 x g for 5 minutes. Cell pellets was re-suspended with 10 ml of mTeSR1 media with 10 uM ROCK inhibitors.

Lipofectamine 3000-DNA transfection complexes were prepared by diluting 10ug of pHAGE-EF1a-BRD4-NUT DNA **(Datafile S1)** or pUC19 (negative control) in 500ul of DMEM/F12 media (ThermoFisher, catalog no. 11330-032), mixing well and incubating for 1 min, followed by addition of 20ul of the P3000 reagent, mixing well and incubating for 1 min at RT. At the same time, 20ul of the Lipofectamine 3000 was diluted in 500ul of DMEM/F12 media, followed by addition of the diluted DNA, mixing well and incubating transfection complexes at RT for 15 minutes. The resulting transfection complexes were added to 7×10^6^ hESC cells in 10ml mTeSR1 media with 10 uM ROCK inhibitors, mixed well and plated cells on a 100mm matrigel coated culture plate. For each immunostaining experiment, matrigel (Corning) coated round glass coverslips were added to the bottom of the plate. 24 hours post-infection, cells were immunostained with antibodies recognizing the NUT protein as described previously.

#### Transfection of mesodermal lineage cells

For mesoderm formation, single hESC cells were plated at a density of approximately 5×10^5^ per well of a 6 well plate (24 wells total) in the presence of ROCK inhibitor and exposed to commercially available mesoderm induction media as instructed by the supplier (Stemcell Technologies 05221). After four days of treatment cells were dissociated into single cells with Accutase (Thermo Fisher) at 37°C for 8 -10 minutes. 7×10^6^ cells were transiently transfected with pHAGE-EF1α-BRD4NUT DNA or pUC19 (negative control) by the same protocol as for hESC except that in all steps, mTeSR1 media (Stemcell Technologies) was replaced with mesoderm induction media (Stemcell Technologies 05221).

### Immunofluorescence

hESC cells were plated in 6 well plates containing matrigel (Corning) coated round glass coverslips. 24 hours post-transfection, cells were immunostained as follows: Coverslips in each well were washed in 3ml 1xPBS at RT, followed by the addition of 2 ml of Fix solution (for 40 ml: 4ml 10xPBS + 24.8ml water+10ml 16% ultra-pure formaldehyde +1.2ml 10% Triton) for 10min. After fixation, coverslips were washed 3x with 3ml of 1xPBS+0.3% Triton X100 for 5 min each, with rocking. 1ml of blocking solution was added per well (for 10ml: 500ul NGS (Normal goat serum) +0.5g BSA+7.5ml water +1ml of 10xPBS+1 ml of 10% Triton X100) and rocked for 1 hour at RT. Blocking solution was then replaced with fresh blocking solution containing anti-NUT antibodies (1:500; Cell Signaling, catalog no. 3625) and incubated overnight at +4C. The next day the coverslips were washed 3x in 3ml of 1xPBS+0.3% Triton X100 for 5 min each, again rocking the 6 well plate. Secondary antibodies (1:1000 anti-Rabbit IgG (H+L) Highly Cross-Adsorbed Secondary Antibody, Alexa Fluor 594, ThermoFisher, catalog no. A-21207) were then added in fresh blocking solution and incubated overnight at +4C. The next day, nuclei were counterstained with ProLong Gold anti-fade reagent with 4’,6-diamidino-2-phenylindole (DAPI) (Life Technologies, catalog no. P36935). Images were viewed on a Zeiss Axioskop 2 using AxioVision Rel. 4.8 software.

### ChIP-seq analysis

We performed ChIP-seq using a rabbit anti-NUT monoclonal antibody (Cell Signaling Technology C52B1). We scaled up the Lipofectamine 3000 transfection protocol to obtain 5×10^7^ hESC and mesodermal lineage cells for each ChIP experiment. BRD4-NUT was expressed for 24 hours prior to harvesting. Two biologically independent experiments were completed for each cell state. The main steps of the ChIP-seq procedure were performed as described in (Alekseyenko et al. 2015a). ChIP-seq reads were aligned to the human reference genome (GRCh37 assembly) using Bowtie (version 2.3.4.3) (Langmead and Salzberg, 2012), retaining only uniquely mapped reads. Smoothed enrichment profiles were generated with a bandwidth of 5 kb. Bigwig files were generated using deeptools (version 3.0.2) with the parameters--RPGC--smoothLength 5000. To visualize ChIP-seq signal at individual regions, we used the Integrative Genomics Viewer (IGV; https://software.broadinstitute.org/software/igv/). Peak enrichment for BRD4-NUT was identified using the HOMER package (Heinz et al. 2010). Briefly, BAM files were converted into tag directories and peaks were called using the findPeaks program with the parameters -region -size 20000 -minDist 200000 -F 2 -L 2 -C 2. 96 kb was used as a minimum size cutoff for megadomain detection. Bedtools intersect (version 1.9) (Quinlan et al. 2010) was used to identify common and differential peaks, and 48 kb was the minimum size scored as an overlap between megadomains in different cell states. Heatmaps for BRD4-NUT occupancy were scaled to an average megadomain size of 180kb summing RPGC-normalized counts from 10 bp bins spanning ± 100 kb from peak start and end site and visualized with the deeptools plotHeatmap function (Ramírez et al. 2014). Heatmaps were plotted to visualize the score distributions of the BRD4-NUT enrichment associated with genomic regions specified in the BED files with peaks from HUES64 or dME cells. Metagene profiles were depicted with the plotProfile function. Genomic coordinates and the identities of genes that map within or flanking each megadomain are listed in **Supp. Datafile S2**.

### Generation of N-BioTAP-cMYC NC797 cells for proteomic analyses

To generate N-BioTAP cMYC fusions in NC797 cells (Toretsky et al. 2003), the following DNA fragment: pAVV-MCS-NheI-5’cMYC_BamHI-ATG-loxP-Blasti-P2A-loxP-PrA-Bio-’cMyc was synthesized as a gBlock (Integrated DNA Technologies, **Datafile S1**) and introduced into the NheI/BamHI restriction sites of pAAV-MCS2 by Gibson assembly (New England Biolabs). The N-BioTAP cassette was targeted to the first ATG of the *MYC* gene. The pAAV-nEFCas9 (Addgene, plasmid 87115) was used to express Cas9, and the pAAV-tagBFP U6-gRNA expression vector was used to express a gRNA targeting the start codon of cMYC: (5’ GACGTTGAGGGGCATCGTCGC ’). Recombinant AAV2 was packaged in HEK293T cells using pHelper and pRC2-mi342 plasmids (Takara, Catalog #632608). Three days after transfection, cells were harvested and AAV2 was isolated using AAVpro Extraction Solution (Takara, Catalog #6235). Two million NC797 cells were infected with 30ul of each adeno-associated virus (AAV2). Following 1 week of culture post-AAV infection, cells were selected with 7.5ug/ml Blasticidin (Invitrogen). Surviving cells were re-plated and single-cell clones were isolated and expanded, and genotyping PCR and sequencing were performed to check for proper integration of the BioTAP cassette in the c-MYC locus. We recovered both heterozygous and homozygous clones. Homozygosity of the cassette insertion was determined by absence of the wildtype PCR fragment (1033bp) and presence of the larger PCR fragment (2083bp) with the following PCR primers: Genom-Myc(1S) 5’ GAGTGGGAACAGCCGCAG ’ and Genom-Myc(1A) 5’ CAGGTACAAGCTGGAGGTGG ’. The presence of the correctly tagged MYC protein was validated by Western blot probed with a c-MYC antibody (Rabbit mAb, Cell Signaling Technology, Cat# 5605). The recombinant pAAV-CRE plasmid was packaged in HEK293T cells using pHelper and pRC2-mi342 plasmids (Takara, Catalog #632608). Cells were harvested three days after transfection and AAV2 was isolated using AAVpro Extraction Solution (Takara, Catalog #6235). Two million 797 cells with homozygous insertions were infected with 30ul of each adeno-associated CRE virus (AAV2). Following one week of culture post-AAV infection, cells were re-plated, single-cell clones were isolated and expanded, and genotyping PCR and sequencing were performed to check for absence of the drug resistance cassette in the c-MYC locus.

### Generation of N-BioTAP-cMYC expressing HEK293T cell for proteomic analyses

We used lentiviral transduction to generate stable HEK293T cell lines (Alekseyenko et al. 2015) carrying an inducible NBioTAP-cMYC transgene. The transgene was constructed using the Gateway recombination system to introduce the Myc ORF clone HsCD00039771 in pDONR221from the CCSB Human ORFeome Collection into the pHAGE-TRE-DEST-NBioTAP (Addgene #53568) lentiviral vector. Stable 293T-Trex-cell lines containing the transgene were generated by lentiviral transduction in the presence of 8µg/mL polybrene, followed by selection with 2µg/mL puromycin dihydrochloride (Sigma #P8833). To induce transcription of the cDNA clones from the CMV/2xtetO promoter we added doxycycline (1µg/mL) to the medium for 48 hours.

### Identification of cMYC interactions using BioTAP-XL

All cell lines were cultured as monolayers in DMEM (Invitrogen) supplemented with 1x Penicillin Streptomycin (Hyclone, South Logan, UT), 1x Glutamax (Gibco), and 10% (v/v) Fetal bovine serum (FBS) (Hyclone). The main steps of the BioTAP-XL procedure were as previously described (Alekseyenko et al. 2017). SDS was removed with ethanol and acetonitrile washes, and proteins were digested with overnight trypsin incubation, yielding peptides for desalting and LC-MS analysis. LC-MS acquisition and analyses were performed as previously described (Alekseyenko et al. 2017). The number of total spectra number for each protein were used to determine relative abundance.

## Supplementary Materials

**Supplemental Figure S1**: IGV browser views of Nascent RNA-seq expression profiles for

*KAT2A, KAT5*, and *TRRAP* in HEK293T and NC797 cells.

**Datafile S1**: DNA sequences of pHAGE-EF1α-BRD4NUT and the Geneblock fragment used to create the N-terminal fusion of BioTAP-to the endogenous *c-MYC* coding region in NC797 cells.

**Datafile S2:** Genomic locations and genes that map within or closely flanking each megadomain.

**Datafile S3**: Total peptide counts for BioTAP-XL analyses of c-MYC in NC797 and HEK293T cells.

## Data Availability

The BioTAP-XL mass spectrometry data are available on the ProteomeXchange Consortium via the PRIDE partner repository, accession number PXD041334. Chip-seq data sets are available on the NCBI GEO database, accession number GSE229558.

## Acknowledgments

We thank Dr. C.A. French for introducing us to NC research, and Ross Tomaino (Taplin Mass Spectrometry Facility, Harvard Medical School) for expert help with the proteomic samples. We thank Drs. T. Martin and S.J. Elledge for the pAAV-tagBFP U6-gRNA expression vector, and Dr. A. Smolko for helpful discussions. This research was funded by the National Institutes of Health, grant number R35 GM126944 to MIK.

## Literature Cited

Alekseyenko AA, Walsh EM, Wang X, Grayson AR, His PT, Kharchenko PV, Kuroda MI, French CA. 2015a. The oncogenic BRD4-NUT chromatin regulator drives aberrant transcription within large topological domains. Genes Dev 29(14): 1507–1523.

Alekseyenko AA, McElroy KA, Kang H, Zee BM, Kharchenko PV, Kuroda MI. 2015b. BioTAP-XL: Cross-linking/tandem affinity purification to study DNA targets, RNA, and protein components of chromatin-associated complexes. Curr Protoc Mol Biol 109: 21.30.1–21.30.32.

Alekseyenko AA, Walsh EM, Zee BM, Pakozdi T, His P, Lemieux ME, Dal Cin P, Ince TA, Kharchenko PV, Kuroda MI, French CA. 2017. Ectopic protein interactions within BRD4-chromatin complexes drive oncogenic megadomain formation in NUT midline carcinoma. Proc Natl Acad Sci U S A 114(21): E4184–E4192.

Bouchard C, Dittrich O, Kiermaier A, Dohmann K, Menkel A, Eilers M, Luscher B. 2001. Regulation of cyclin D2 gene expression by the Myc/Max/Mad network: Myc-dependent TRRAP recruitment and histone acetylation at the cyclin D2 promoter. Genes Dev 15(16): 2042–2047.

Chen AE, Egli D, Niakan K, Deng J, Akutsu H, Yamaki M, Cowan C, Fitz-Gerald C, Zhang K, Melton DA, Eggan K. 2009. Optimal timing of inner cell mass isolation increases the efficiency of human embryonic stem cell derivation and allows generation of sibling cell lines. Cell Stem Cell 4(2): 103–106.

Dang CV, Reddy EP, Shokat KM, Soucek L. 2017. Drugging the ‘undruggable’ cancer targets. Nat Rev Cancer 17(8): 502–508.

Dingar D, Kalkat M, Chan PK, Srikumar T, Bailey SD, Tu WB, Coyaud E, Ponzielli R, Kolyar M, Jurisica I, Huang A, Lupien M, Penn LZ, Raught B. 2015. BioID identifies novel c-MYC interacting partners in cultured cells and xenograft tumors. J Proteomics 118: 95–111.

Eagen KP, French CA 2021. Supercharging BRD4 with NUT in carcinoma. Oncogene 40(8): 1396–1408.

Facchini LM, Penn LZ. 1998. The molecular role of Myc in growth and transformation: recent discoveries lead to new insights. FASEB J 12(9): 633–651.

Filippakopoulos P, Qi J, Picaud S, Shen Y, Smith WB, Fedorov O, Morse EM, Keates T, Hickman TT, Felletar I, Philpott M, Munro S, McKeown MR, Wang Y, Christie AL, West N, Cameron MJ, Schwartz B, Heightman TD, La Thangue N, French CA, Wiest O, Kung AL, Knapp S, Bradner JE. 2010. Selective inhibition of BET bromodomains. Nature 468(7327): 1067–1073.

Frank SR, Parisi T, Taubert S, Fernandez P, Fuchs M, Chan HM, Livingston DM, Amati B. 2003. MYC recruits the TIP60 histone acetyltransferase complex to chromatin. EMBO Rep 4(6): 575–580.

Frank SR, Schroeder M, Fernandez P, Taubert S, Amati B. 2001. Binding of c-Myc to chromatin mediates mitogen-induced acetylation of histone H4 and gene activation. Genes Dev 15(16): 2069–2082.

French CA. 2018. NUT Carcinoma: Clinicopathologic features, pathogenesis, and treatment. Pathol Int 68(11): 583–595.

French CA, Miyoshi I, Kubonishi I, Grier HE, Perez-Atayde AR, Fletcher JA. 2003. BRD4-NUT fusion oncogene: a novel mechanism in aggressive carcinoma. Cancer Res 63(2): 304–307.

Grayson AR, Walsh EM, Cameron MJ, Godec J, Ashworth T, Ambrose JM, Aserlind AB, Wang H, Evan G, Kluk MJ, Bradner JE, Aster JC, French CA. 2014. MYC, a downstream target of BRD-NUT, is necessary and sufficient for the blockade of differentiation in NUT midline carcinoma. Oncogene 33(13): 1736–1742.

Heinz S, Benner C, Spann N, Bertolino E, Lin YC, Laslo P, Cheng JX, Murre C, Singh H, Glass CK. 2010. Simple combinations of lineage-determining transcription factors prime cis-regulatory elements required for macrophage and B cell identities. Molecular Cell, 38(4), 576–589.

Henriksson M, Luscher B. 1996. Proteins of the Myc network: essential regulators of cell growth and differentiation. Adv Cancer Res 68: 109–182.

Ibrahim Z, Wang T, Destaing O, Salvi N, Hoghoughi N, Chabert C, Rusu A, Gao J, Feletto L, Reynoird N, Schalch T, Zhao Y, Blackledge M, Khochbin S, Panne D. 2022. Structural insights into p300 regulation and acetylation-dependent genome organisation. Nat Commun 13(1): 7759.

Kalkat M, Resetca D, Lourenco C, Chan PK, Wei Y, Shiah YJ, Vitkin N, Tong Y, Sunnerhagen M, Done SJ, Boutros PC, Raught B, Penn LZ. 2018. MYC Protein interactome profiling reveals functionally distinct regions that cooperate to drive tumorigenesis. Mol Cell 72(5): 836–848 e837.

Kim J, Woo AJ, Chu J, Snow JW, Fujiwara Y, Kim CG, Cantor AB, Orkin SH. 2010. A Myc network accounts for similarities between embryonic stem and cancer cell transcription programs. Cell 143(2): 313–324.

Kotekar A, Singh AK, Devaiah BN. 2022. BRD4 and MYC: power couple in transcription and disease. FEBS J.

Langmead B, Salzberg SL. 2012. Fast gapped-read alignment with Bowtie 2. Nature methods, 9(4), 357–359.

Lee VM, Andrews PW. 1986. Differentiation of NTERA-2 clonal human embryonal carcinoma cells into neurons involves the induction of all three neurofilament proteins. J Neurosci 6(2): 514–521.

Liao S, Maertens O, Cichowski K, Elledge SJ. 2018. Genetic modifiers of the BRD4-NUT dependency of NUT midline carcinoma uncovers a synergism between BETis and CDK4/6is. Genes Dev 32(17-18): 1188–1200.

Liu X, Tesfai J, Evrard YA, Dent SY, Martinez E. 2003. c-Myc transformation domain recruits the human STAGA complex and requires TRRAP and GCN5 acetylase activity for transcription activation. J Biol Chem 278(22): 20405–20412.

Malynn BA, de Alboran IM, O’Hagan RC, Bronson R, Davidson L, DePinho RA, Alt FW. 2000. N-myc can functionally replace c-myc in murine development, cellular growth, and differentiation. Genes Dev 14(11): 1390–1399.

McMahon SB, Wood MA, Cole MD. 2000. The essential cofactor TRRAP recruits the histone acetyltransferase hGCN5 to c-Myc. Mol Cell Biol 20(2): 556–562.

Mustachio LM, Roszik J, Farria A, Dent SYR. 2020. Targeting the SAGA and ATAC transcriptional coactivator complexes in MYC-driven cancers. Cancer Res 80(10): 1905–1911.

Naxerova K, Di Stefano B, Makofske JL, Watson EV, de Kort MA, Martin TD, Dezfulian M, Ricken D, Wooten EC, Kuroda MI, Hochedlinger K, Elledge SJ. 2021. Integrated loss- and gain-of-function screens define a core network governing human embryonic stem cell behavior. Genes Dev 35(21-22): 1527–1547.

Quinlan AR, Hall IM. 2010. BEDTools: a flexible suite of utilities for comparing genomic features. Bioinformatics (Oxford, England), 26(6), 841–842.

Ramírez F, Dündar F, Diehl S, Grüning BA, Manke T. 2014. deepTools: a flexible platform for exploring deep-sequencing data. Nucleic acids research, 42(Web Server issue), W187–W191.

Rousseaux S, Reynoird N, Khochbin S. 2022. NUT is a driver of p300-mediated histone hyperacetylation: from spermatogenesis to cancer. Cancers (Basel) 14(9).

Roux KJ, Kim DI, Burke B, May DG. 2018. BioID: A Screen for Protein-Protein Interactions. Curr Protoc Protein Sci 91: 19 23 11–19 23 15.

Shiota H, Barral S, Buchou T, Tan M, Coute Y, Charbonnier G, Reynoird N, Boussouar F, Gerard M, Zhu M, Bargier L, Puthier D, Chuffart F, Bourova-Flin E, Picaud S, Filippakopoulos P, Goudarzi A, Ibrahim Z, Panne D, Rousseaux S, Zhao Y, Khochbin S. 2018. Nut directs p300dependent, genome-wide H4 hyperacetylation in male germ cells.舡 Cell Rep 24(13): 3477–3487 e3476.

Toretsky JA, Jenson J, Sun CC, Eskenazi AE, Campbell A, Hunger SP, Caires A, Frantz C, Hill JL, Stamberg J. 2003. Translocation (11;15;19): a highly specific chromosome rearrangement associated with poorly differentiated thymic carcinoma in young patients. Am J Clin Oncol 26(3): 300–306.

Wang R, You J. 2015. Mechanistic analysis of the role of bromodomain-containing protein 4 (BRD4) in BRD4-NUT oncoprotein-induced transcriptional activation. J Biol Chem 290(5): 2744–2758.

Yu D, Liang Y, Kim C, Jaganathan A, Ji D, Han X, Yang X, Jia Y, Gu R, Wang C, Zhang Q, Cheung KL, Zhou MM, Zeng L. 2023. Structural mechanism of BRD4-NUT and p300 bipartite interaction in propagating aberrant gene transcription in chromatin in NUT carcinoma. Nat Commun 14(1): 378.

Zeng L, Zhou MM. 2002. Bromodomain: an acetyl-lysine binding domain. FEBS Lett 513(1): 124–128.

